# Number neurons in the nidopallium of young domestic chicks

**DOI:** 10.1101/2022.01.21.475044

**Authors:** Dmitry Kobylkov, Uwe Mayer, Mirko Zanon, Giorgio Vallortigara

## Abstract

Numerical cognition is ubiquitous in the animal kingdom. Domestic chicks are a widely used developmental model for studying numerical cognition. Soon after hatching, chicks can perform sophisticated numerical tasks. Nevertheless, the neural basis of their numerical abilities has remained unknown. Here, we describe for the first time number neurons in the caudal nidopallium (functionally equivalent to the mammalian prefrontal cortex) of young domestic chicks. Number neurons that we found in young chicks showed remarkable similarities to those in the prefrontal cortex and caudal nidopallium of adult animals. Thus, our results suggest that numerosity perception based on the labeled-line code provided by number neurons might be an inborn feature of the vertebrate brain.

**Significance:** Numerosity, i.e. the number of items in a set, is a significant aspect in the perception of the environment. Behavioural and in silico experiments suggest that number sense belongs to a core knowledge system and can be present already at birth. However, neurons sensitive to the number of visual items have been so far described only in the brain of adult animals. Therefore, it remained unknown to what extent their selectivity would depend on visual learning and experience. We found number neurons in the caudal nidopallium (a higher associative area with functional similarities to the mammalian prefrontal cortex) of very young, numerically naïve domestic chicks. This result suggests that numerosity perception is possibly an inborn feature of the vertebrate brain.

## Introduction

Be it a number of conspecifics in a group (Balestrieri et al. 2019), a number of food items (Hunt et al. 2008), or a number of motifs in a song (Templeton et al. 2005), correct estimation of quantities is of vital importance for animals. Several behavioural studies have confirmed that numerical competence is not a prerogative of human beings, but is a widespread phenomenon in the animal kingdom (reviewed by Nieder 2019, Lorenzi et al. 2021). Mammals (Davis and Albert 1986, Ward and Smuts 2007, Beran et al. 2008), birds (Lyon 2003, Templeton et al. 2005, Rugani et al. 2009), reptilians (Gazzola et al. 2018), amphibians (Stancher et al. 2015), fishes (Potrich et al. 2015), and invertebrates (Bortot et al. 2021), although evolutionary distant, all can spontaneously assess quantities using an approximate number system (Brannon and Merritt 2011).

For the approximate number system, which is based on the Weber-Fechner law (Nieder 2016), the perception of cardinal numbers resembles the perception of continuous physical stimuli. As a consequence, discrimination of quantities is imprecise and depends on the numerical distance between stimuli. In other words, it is easier to tell apart 5 and 10 than 9 and 10. Moreover, discrimination of quantities becomes increasingly difficult with the numerical size. For a given numerical distance (e.g., 1) it is easier to discriminate between numbers with low magnitudes (1 vs. 2), than with high magnitudes (9 vs. 10).

Recent research has uncovered that the approximate number system relies on the activity of a specific neuronal population. Neurons that respond to abstract numerosity irrespective of objects’ physical appearance (shape, colour, size) have been found in the forebrain of human and non-human primates (Nieder 2012, Kutter et al. 2018) and in crows (Ditz and Nieder 2015). In mammals, numerical responses were recorded in the parietal and the prefrontal cortices (PFC, Nieder 2012). In birds, similar neurons have been described in the caudolateral nidopallium (NCL, Ditz and Nieder 2015). The NCL is believed to be an analogue of the PFC in the avian brain (Güntürkün et al. 2021) and is involved in a variety of cognitive processes, including memory formation (Diekamp et al. 2002, Hahn et al. 2021), abstract rule learning (Veit and Nieder 2013), and action planning (Veit et al. 2015).

Both monkeys and crows are among the most evolutionary advanced species of their phylogenetic groups. They independently developed sophisticated intellectual capacities (Nieder et al. 2020) and both possess enlarged forebrains (Olkowicz et al., 2016). The neural representation of numerosities described in these species also shares remarkable similarities (Nieder et al. 2002, Ditz and Nieder 2015, Viswanathan and Nieder 2013, Wagener et al. 2018). In both species, the number neurons tuned to a preferred numerosity show a gradual decrease in firing rate to other numerosities (numerical distance effect). Their tuning curves are skewed towards larger numerosities and become progressively broader (less selective) with increasing numerosities (numerical size effect). However, it is unclear whether the presence of similar number neurons in these two species emerges as a consequence of their elaborate cognitive skills and enlarged forebrains. To understand the evolution of the number sense we need to explore its neural correlates in distant bird species with more ancestral traits.

Moreover, until now, number neurons have been described only in adult animals (e.g. Nieder et al. 2002, Sawamura et al. 2002, Viswanathan and Nieder 2013, Ditz and Nieder 2015, Wagener et al. 2018). At the same time, behavioural data from human infants (Izard et al. 2009) and young domestic chicks (Rugani et al. 2008, 2009) indicate that some core numerical abilities might be an inborn or spontaneously emerging (Kim et al. 2021, Nasr et al. 2019) property of the vertebrate brain. Testing the presence of number neurons in young and untrained organisms is crucial to verify this hypothesis.

In our study, we aimed to describe the neural correlates of the number sense in domestic chicks (*Gallus gallus*), which belong to a sister group of modern Neoaves (Prum et al. 2015). The domestic chick is a well-established developmental model for studying numerical cognition. Soon after hatching these birds are already capable of discriminating quantities (Rugani et al. 2008, 2013) and even performing basic arithmetic operations (Rugani et al. 2009). It has also been shown that young chicks represent numbers across the mental number line (Rugani et al. 2015), a cognitive ability that had been previously attributed only to humans.

We hypothesised that neural processing of numerical information in young untrained chicks might be similar to crows, despite them having been evolving independently over the last ∼ 70 million years (Prum et al. 2015). In a domestic chicken, the NCL is morphologically different from that of corvids (von Eugen et al. 2020), but it is unclear whether this reflects any functional difference. Therefore, we decided to search for neural responses to numerical stimuli in the NCL of domestic chicks. For this purpose we habituated young chicks to a computer monitor, where numerical stimuli were presented (Fig. 1A). We explored neural responses to numerosities from 1 to 5. To control for non-numerical parameters we presented three different categories of stimuli:“radius-fixed”, “area-fixed”, and “perimeter-fixed” (Fig. 1B).

**Figure 1.**
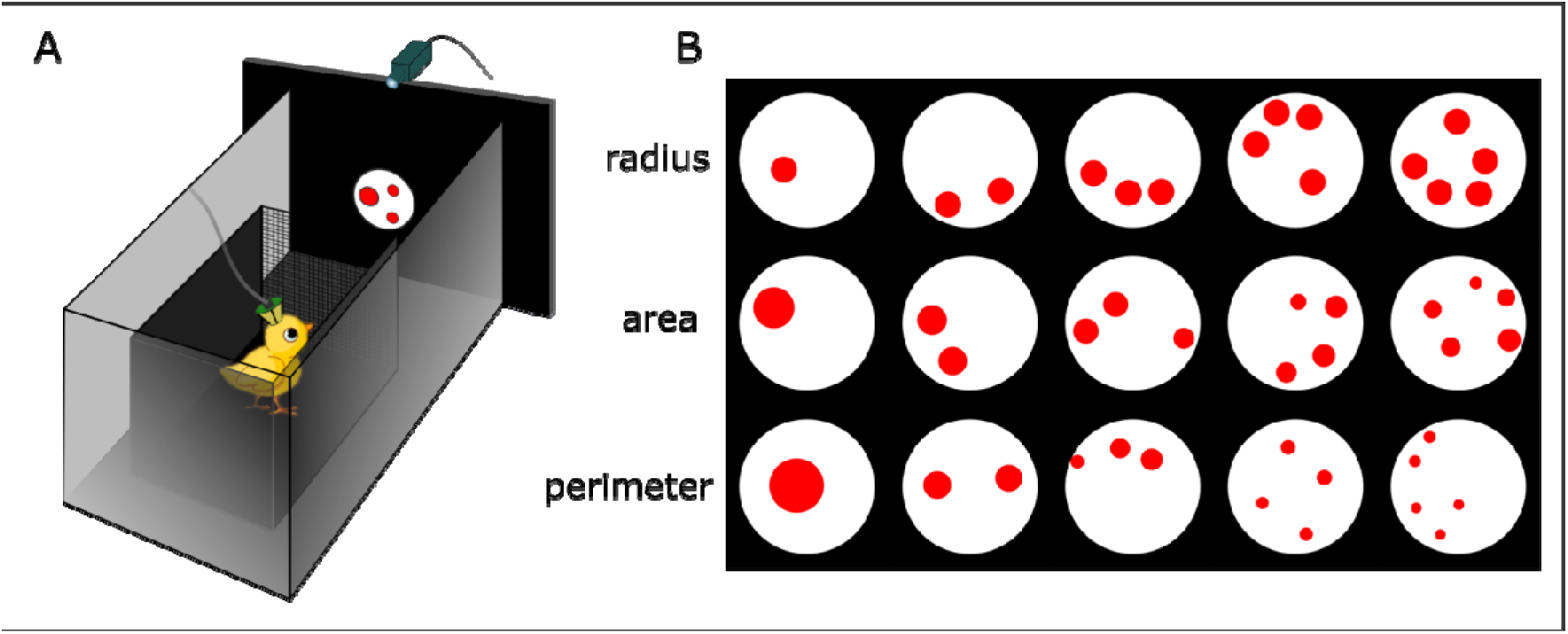
Experimental design. (A) Schematic drawing of the experimental setup. Young chicks were placed in a small wooden box in front of the screen, where numerical stimuli appeared. They were trained to pay attention to the stimuli without any further discrimination between different numerosities. (B) Examples of different types of numerosity stimuli that we presented by every neural recording: “radius-fixed”, “area-fixed”, and “perimeter-fixed”.

## Results

We recorded the activity of 471 units in the NCL of young domestic chicks and examined the mean firing rate of each unit during stimulus presentation (Fig. 2A-B). Among these units, 53 (11%) responded to numerosity irrespective of the stimulus type (henceforth “number neurons”, see Nieder 2016). This was revealed by a two-way ANOVA with the factors “numerosity” (5 levels: 1-5) and “stimulus type” (3 levels: “radius-fixed”, “area-fixed”, “perimeter-fixed”). The unit was considered as a number neuron only if the main effect of the factor “numerosity” was highly significant (p<0.01), but not the effect of the “stimulus type” or the interaction between the two factors. The numerosity that elicited the strongest neural response was defined as the preferred numerosity for this number neuron (after Ditz and Nieder 2015). Five examples of number neurons tuned to different numerosities are shown in Fig. 2C-G. The corresponding statistics are summarised in Table 1. The responses of all 53 number neurons are summarised in Fig. 3A (for statistical results see Table S1 in the supplementary materials, for an example of trials with numerical responses see Video S1 in the supplementary materials).

**Figure 2.**
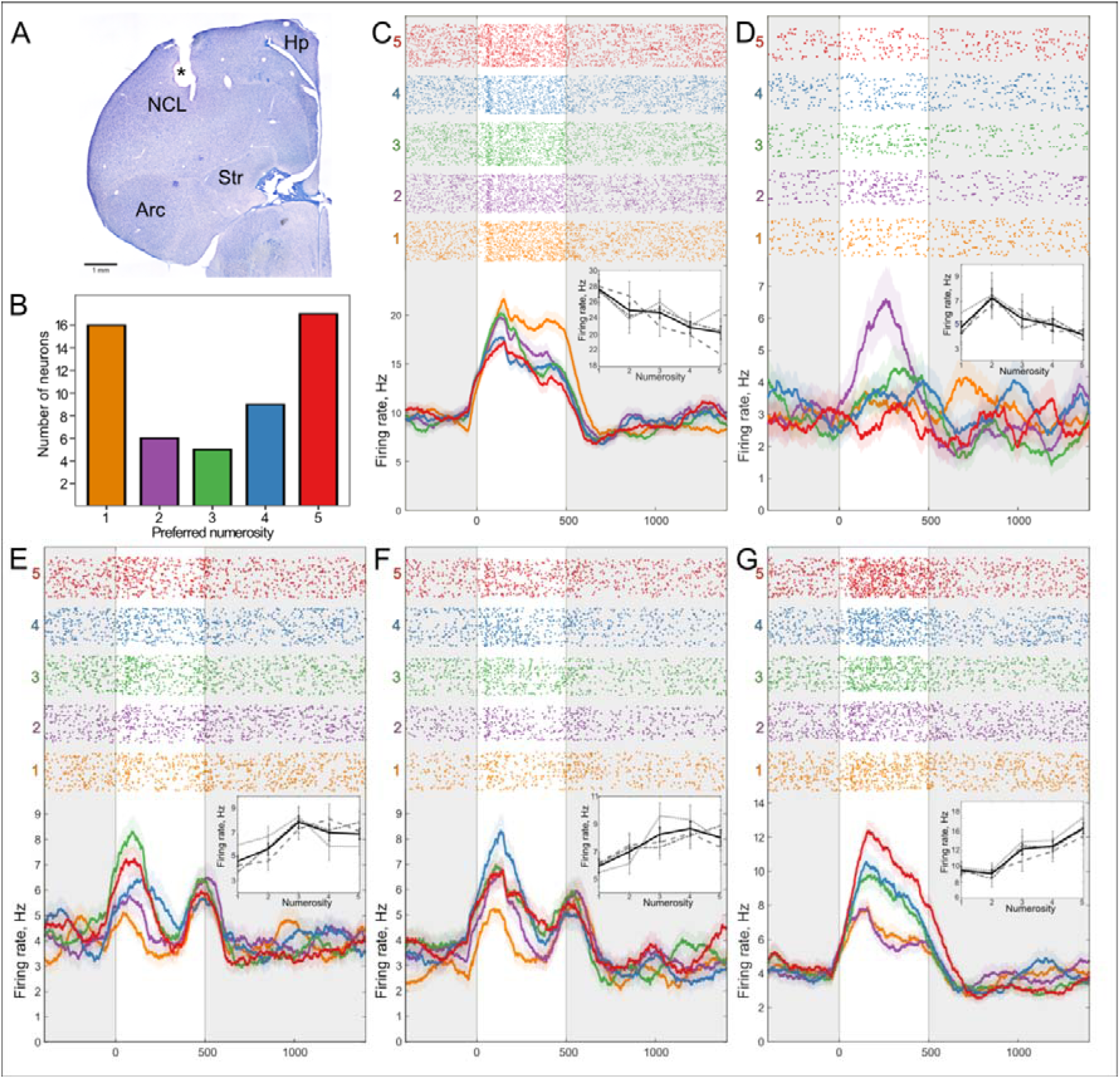
Neurons in the NCL of chicks responding to numerosity. (A) An exemplary coronal section of the chicken forebrain showing the recording site in the NCL (electrolytic lesion is marked by an asterisk). Arc: arcopallium, Hp: hippocampus, NCL: caudolateral nidopallium, Str: striatum. (B) Distribution of neurons that preferred each numerosity stimulus. Examples of neurons that were tuned to numerosity 1 (C), 2 (D), 3 (E), 4 (F), or 5 (G). Top: raster plots representing neural activity, where each line corresponds to one trial and each dot corresponds to a spike. Trials are grouped by numerosity. The 500-ms duration of the stimulus is marked by a transparent window. Bottom: averaged spike density functions (smoothed by a 100 ms Gaussian kernel; SEM is plotted as a shaded area along the lines). Insert: average firing rate in response to numerosities of each stimulus type. Grey dotted line corresponds to “radius-fixed”, dashed line – to “perimeter-fixed”, dot-dashed line – to “area-fixed” stimuli, and black solid line – to an average. Error bars correspond to SEM.

**Figure 3.**
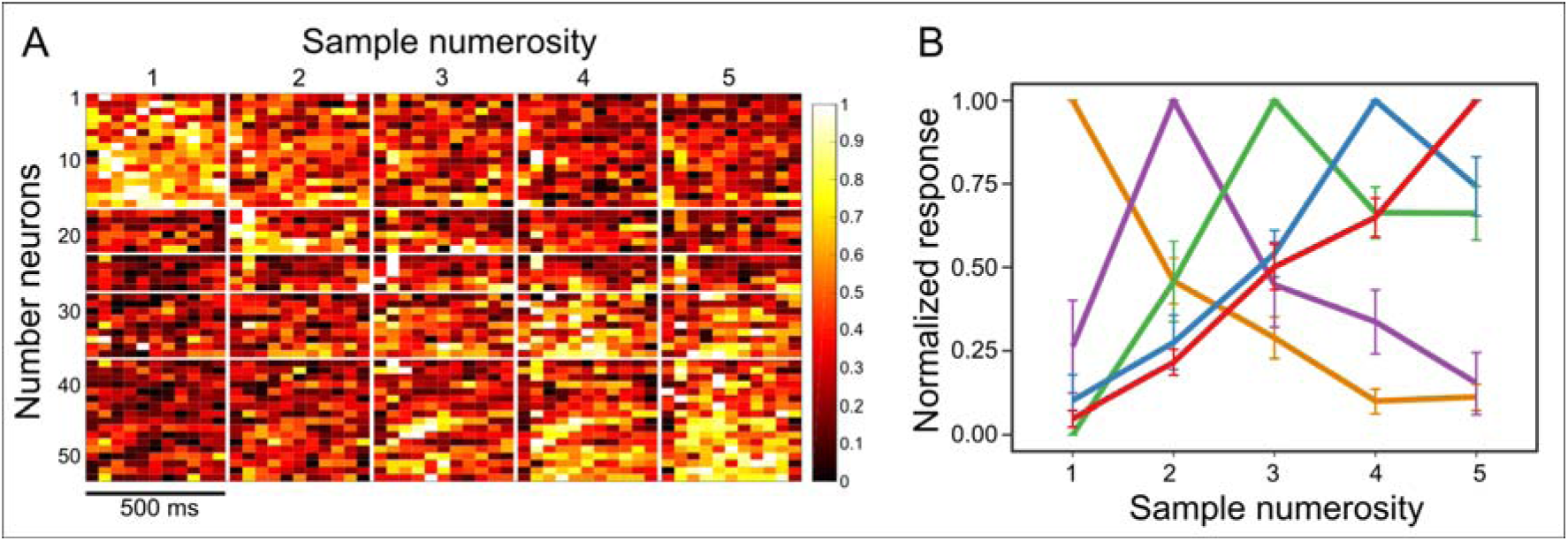
(A) Neural response of all recorded number neurons to numerosity stimuli. Heatmap values represent the mean firing rate during the stimulus presentation (binned by 50-ms), normalised [0, 1] for the corresponding neuron in each row. Values are further grouped by the numerosity stimuli from 1 to 5 (vertical white lines), and by the numerical preference of recorded neurons (horizontal white lines) from neurons that preferred numerosity 1 (top) to neurons that preferred numerosity 5 (bottom). (B) Average tuning curves of numerosity selective neurons. The neural activity of neurons is first normalised [0 = response to the least-preferred numerosity, 1 = response to the most-preferred numerosity] and then grouped by their most preferred numerosity. Error bars correspond to SEM.

**Table 1.** Results of the two-way ANOVA for five example number neurons shown in the Figure 2 (C-G). Preferred numerosity: numerosity eliciting the strongest response. ANOVA results (F-statistics and p-value) for the factor “stimulus type” (“radius-fixed”, “area-fixed”, “perimeter-fixed”), “numerosity” (numerosity 1 to 5), or interaction between them.

| Preferred | ANOVA (Firing rate ~ Stimulus type * Numerosity) |  |  |
| --- | --- | --- | --- |
| numerosity | Numerosity | Stimulus type | Interaction |
| num1 | F(4,251)=5.7273, p < 0.001 | F(2,251)=0.9275, p = 0.396 | F(8,251)=1.2411, p = 0.273 |
| num2 | F(4,253)=3.5881, p = 0.007 | F(2,253)=0.2355, p = 0.79 | F(8,253)=0.454, p = 0.887 |
| num3 | F(4,496)=5.8789, p < 0.001 | F(2,496)=0.1179, p = 0.889 | F(8,496)=1.4538, p = 0.173 |
| num4 | F(4,308)=4.2385, p = 0.002 | F(2,308)=0.1141, p = 0.892 | F(8,308)=0.9124, p = 0.506 |
| num5 | F(4,355)=13.4938, p < 0.001 | F(2,355)=1.832, p = 0.162 | F(8,355)=0.3219, p = 0.958 |

Most of the number neurons were tuned to the numerosity 1 (30%, N = 16) and 5 (32%, N = 17). However, we found neurons responsive to other numerosities as well (2: 11%, N = 6; 3: 9%, N = 5; 4: 17%, N = 9), covering the whole range of tested numerosities (Fig. 2B).

When grouped by their preferred numerosity, the tuning curves of number neurons were asymmetric and increasingly wider towards larger numerosities (Fig. 3B). To quantify this numerical magnitude effect, we plotted neural filter functions of single neurons on four different scales: a linear scale, a power function with an exponent of 0.5, a power function with an exponent of 0.33, and a logarithmic (log2) scale. The neural filter functions became significantly more symmetric on a non-linear scale (Friedman test: X^2^(3) = 10.653; p = 0.014, Fig. 4A), with the linear scale significantly different from the power 0.33 and logarithmic scales (Nemenyi test: p = 0.022 and p = 0.028, respectively), but not from the power 0.5 scale (Nemenyi test: p = 0.176). The sigma of the Gaussian fit increased with numerosity only when plotted on a linear scale (Fig. 4B, slope of the linear fit = 0.17) but not with other non-linear scales (Fig. 4B, power 0.5: slope = -0.003, power 0.33: slope = -0.01, log2: slope = -0.03).

**Figure 4.**
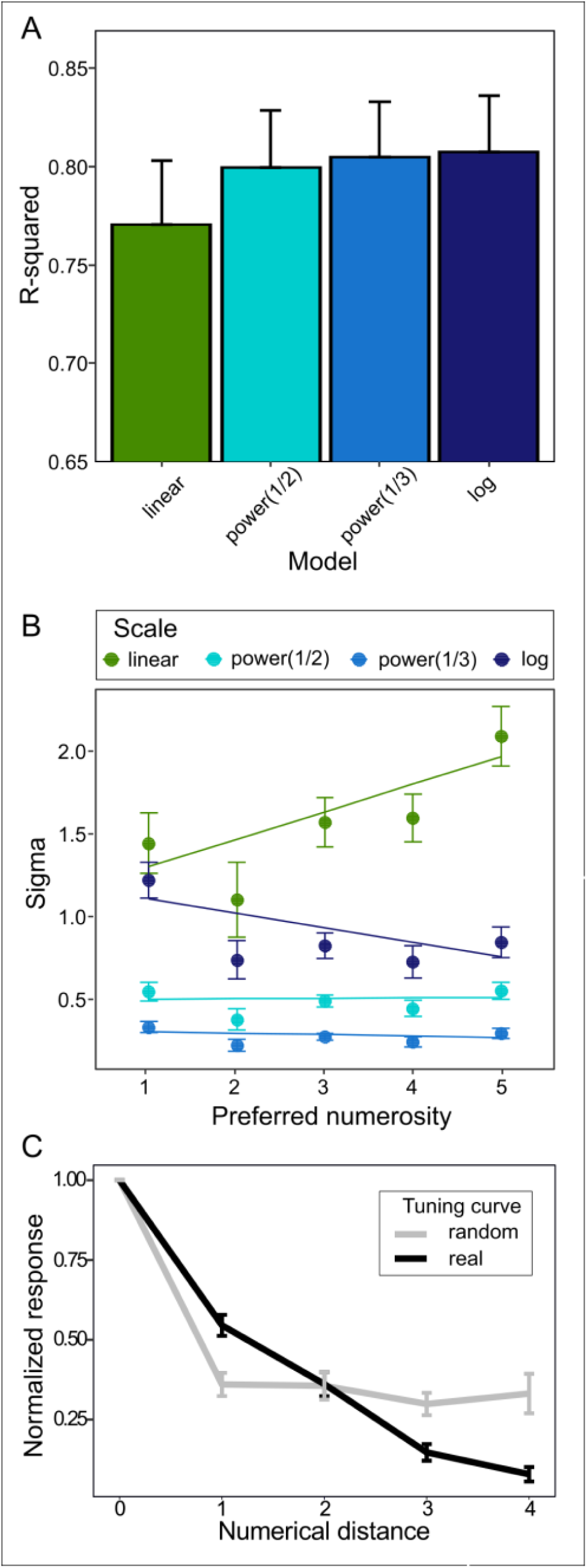
(A-B) Comparison of different scaling schemes for the tuning curves. (A) R-squared, a measure of goodness-of-fit reflects symmetry of the tuning curves plotted on the four different scales. The tuning curves of number neurons become more symmetric when plotted on the non-linear scale. (B) Sigma of the Gaussian fit for neurons preferring different numerosities. When plotted on the linear scale, the tuning curves become wider with increased numerosity. Error bars correspond to SEM. (C) Averaged normalised activity of all numerosity selective neurons compared to the random tuning curve (see Materials and methods section for details). The neural activity was normalised [0 = response to the least-preferred numerosity, 1 = response to the most-preferred numerosity] and then plotted as a function of absolute numerical distance from the most preferred numerosity. Neural response of numerosity selective neurons (black line) gradually decreased with the numerical distance. The slope of this tuning curve is notably different from the random tuning curve (grey line) of false-positive neurons obtained by random shuffling of trials. Error bars correspond to SEM.

To evaluate the probability of finding false positive numerical responses in our dataset, we used the trial-shuffle method (1000 times for every recorded unit). After each shuffling we selected false-positive number neurons by the same statistical criteria described above. The proportion of real number neurons (11 %; 53 out of 471) was significantly higher than the proportion of false-positive units (0,95%; 4488 out of 471000) obtained by the analysis of randomly shuffled trials (proportion test: X^2^(1) = 512,56, p<0.001).

We further compared the tuning curves between the false positive number neurons and the real number neurons. For this, for each numerical category we randomly selected the same number of corresponding false-positive neurons as the actually recorded number neurons in that category (Fig 2B). We compared the tuning curves of the real and false-positive neurons (Fig 4C) computing a two-way ANOVA with the factors “absolute numerical distance” (5 levels: 0-4) and “data type” (2 levels: real/false positive). The response of real neurons decreased gradually with numerical distance, meaning that closest numerosities were more likely to trigger the number neurons. In contrast, the tuning curve of the false-positive neurons, which by chance happened to have higher firing rates for a random numerical stimulus, was markedly different from real number neurons (interaction for the factor “numerical distance * data type”: F_(1,526)_ = 6.275; p = 0.01).

We further described the selectivity of number neurons for unique numerosities by performing a post hoc analysis and comparing the response between the most preferred and other numerosities (Table 2). 9 out of 53 number neurons showed statistically different firing rates between the most-preferred and closest neighbouring numerosities. As expected from the numerical size effect, neurons tuned to lower numerosities are generally more selective than neurons coding for larger numerosities. 6 out of 16 neurons with the preferred numerosity 1 significantly decreased their firing rate in response to the numerosity 2. At the same time, out of 17 neurons that preferentially responded to the numerosity 5, only 1 neuron showed significantly lower response to the numerosity 4. The number neurons generally better differentiated between numerosities with increasing numerical distance (GLM for absolute numerical distance: X^2^(18) = 3.935, p = 0.047; GLM for the log of the numerical distance: X^2^(18) = 6.544, p = 0.011). This effect was more prominent when the numerical distance was calculated on the logarithmic scale: the GLM for the log of the numerical distance had a better fit than the GLM for absolute numerical distance (ΔBIC = 6.36). For the summary of all post hoc results, see TableS2 in the supplementary materials.

**Table 2.** The summary of the post hoc analysis. For each group based on their preferred numerosity we calculated the number of neurons that showed significant difference between the most-preferred and the given numerosity.

| Summary |  | Number of neurons |  |  |  |  |
| --- | --- | --- | --- | --- | --- | --- |
| Sum | Group | num1 | num2 | num3 | num4 | num5 |
| 16 | num1 |  | 6 | 9 | 12 | 14 |
| 6 | num2 | 4 |  | 2 | 1 | 3 |
| 5 | num3 | 5 | 1 |  | 0 | 0 |
| 9 | num4 | 6 | 2 | 1 |  | 1 |
| 17 | num5 | 15 | 12 | 6 | 1 |  |

## Discussion

We recorded number neurons in 8-12 day old chicks, the youngest animals in which number neurons have been described so far. In the NCL of young domestic chicks, 11% of neurons showed a strong selective response to numerical stimuli, confirming the role of this structure in avian numerical cognition. Moreover, our chicks were not trained in any numerical discrimination task. Instead, they were simply paying attention to the screen, where the numerical stimuli were presented (passive fixation, Hussar and Pasternak 2009). Thus, our young chicks can be considered numerically naïve (see Viswanathan and Nieder 2013).

This result is in line with previous studies on numerically naïve adult animals of distant lineages: 14% of neurons in the prefrontal cortex (PFC) of adult monkeys (Viswanathan and Nieder 2013) and 12% of neurons in the NCL of adult crows (Wagener et al. 2018) have shown numerical response. It is important to note that in any of these studies including our own, we cannot completely exclude a potential unsupervised learning effect due to a repeated exposure to numerical stimuli. Indeed, recent studies have found that in a deep neural network designed to analyse images, a similar amount of number detectors can emerge spontaneously (Nasr et al. 2019), even without pre-training (Kim et al. 2021). However, the hypothesis that number neurons can spontaneously emerge in the biological visual system has not been directly tested so far.

Our data, together with behavioural evidence from newly hatched domestic chicks (Rugani et al. 2008, 2009) and newborn infants (Izard et al. 2009), suggest that numerosity detection might be an inborn or spontaneously emerging feature of the brain. Our results certainly should not be automatically extrapolated to other newborn organisms, since the domestic chicken is a precocial species with a very rapid development after hatching. Nevertheless, even in 2-weeks-old chicks, the functional organization of the brain still remains immature and flexible (Zappia and Rogers 1987). Thus, the presence of number neurons in young chicks, which were not trained to discriminate any specific numerosity, supports the idea of an inborn number sense. Moreover, since behavioural studies (Rugani et al. 2009, 2013) show that newly hatched chicks perceive numerical information, we might expect to find number neurons already at the first day after hatching.

The number neurons in domestic chicks showed very similar features to those observed in primates and crows (Viswanathan and Nieder 2013, Ditz and Nieder 2015, 2016). First, the number neurons we observed were tuned to specific numerosities, in accordance with the labelled-line code shown for the number neurons in the NCL of crows (Ditz and Nieder 2015, 2016). Second, chicks’ number neurons showed a specific decay in response to non-preferred numerosities, which was similar to number neurons in other species and could not have been obtained by chance (see false-positive neurons in Fig. 4C). Third, we observed the numerical magnitude effect: the tuning curves of number neurons became wider with increased numerosity and more symmetrical when plotted on the non-linear scale (Fig. 3B). Also the selectivity of the neural response to non-preferred numerosities increased with the logarithm of the numerical distance, rather than with the absolute numerical distance.

The overall low selectivity to specific numerosity at a single-cell level, as revealed by the post hoc analysis (Table 2), appears to be similar to what was observed in other species (Viswanathan and Nieder 2013; Ditz and Nieder 2015, 2016). How brains can depict precise numerosities with such a noisy system may be explained by a population rate code (Nieder 2016). Single cells respond to every trial in a probabilistic way and only on average show increased firing rates to a given numerosity. Smooth tuning curves only emerge after the neural filter functions of many single cells tuned to the same numerosity are pooled together (Fig. 3B). For an animal to immediately assess numerosity, independent responses of several number neurons should be averaged simultaneously over a large population. Thus, at the neural population level, stimulation by a specific numerosity would result in a distinct activation pattern. This hypothesis can be tested only by implementing methods that allow recording of single-cell activity over a large neural population.

Given these striking similarities between number neurons in the NCL of crows and chicks, it seems a reasonable guess that they share the same evolutionary origin. This, in turn, would mean that the neural mechanism of the number sense is not an advanced evolutionary adaptation of a few highly intelligent species. Number neurons in the NCL are likely an ancestral feature in avian species, since domestic chicks belong to the sister group of modern Neoaves (Prum et al. 2015).

While it is very tempting to go one step further and discuss the idea of the common evolutionary origin of number neurons for all vertebrates, this would be too speculative. The NCL in birds and the prefrontal cortex in mammals do share similar functions, but are not homologous structures (Güntürkün and Bugnyar 2016, Preuss and Wise 2021). Birds independently developed cortical brain regions, including the NCL, that enabled their high cognitive functions (Güntürkün et al. 2021) based on similar to mammalian pallial architecture (Stacho et al. 2020). The same is likely true also for the Dc (dorsal-central) region in the telencephalon of zebrafish, which has been recently shown to process numerical information (Messina et al. 2021). The possibility of a direct homology between the Dc and either the PFC or the NCL can only be addressed at a macroscopic level. These three regions likely belong to larger neocortex homologues in the respective species (Briscoe and Ragsdale 2019, but see Puelles 2017). However, given the developmental and morphological differences between these areas, new cortical subregions with similar functions must have appeared multiple times during independent evolution of these structures.

Two hypotheses can be put forward on the evolution of number sense in vertebrates. The first one is that number perception evolved independently several times in different phylogenetic groups, although our data strongly suggest that number neurons in the nidopallium are an ancestral trait of birds. This mechanism, however, still can be an adaptation that evolved in parallel to mammalian number neurons. In this case, the identical labelled-line coding scheme for numerosities adopted by birds and mammals might be computationally advantageous and, therefore, evolved independently in both groups (Ditz and Nieder 2015). The second hypothesis is that numerosity processing can be based upon an ancient core neural circuit shared among all vertebrates. In this case, we would expect to find numerosity responses in other, evolutionary conserved brain regions homologically shared among vertebrates (Lorenzi et al. 2021). Indeed, some indirect evidence suggests that at least a coarse estimation of quantity, i.e. more vs. less, might be present already at the subcortical level in humans (Collins et al. 2017) and in the midbrain of birds (Gusel’nikov et al. 1971) and zebrafish (Preuss et al. 2014).

These hypotheses are, however, not mutually exclusive. The putative ancestral neural circuit might be dedicated to assess continuous physical parameters normally associated with the numerosity, like total area of the stimulus. On the contrary, higher order brain processing leading to estimation of cardinal numerosities at a more abstract level may have developed several times, together with the independent evolution of new cortex homologue brain areas in distant phylogenetic groups (Striedter and Northcutt 2019).

Summing up, our study provides the first step in addressing a complex evolutionary and developmental aspect of numerical cognition. By implementing a simplified training procedure, we could, for the first time, demonstrate the existence of number neurons in young numerically naïve domestic chicks. In the future, this method might be easily adopted for studying the neural correlates of numerical cognition in other brain regions, as well as in other species.

## Materials and methods

### Subjects

12 domestic chicks (*Gallus gallus domesticus*) of both sexes from the Aviagen ROSS 308 strain were used. Fertilized eggs were obtained from a local commercial hatchery (CRESCENTI Società Agricola S.r.l.–AllevamentoTrepola–cod. Allevamento 127BS105/2). Eggs were incubated and hatched within incubators (Marans P140TU-P210TU) at a temperature of 37.7 °C, with 60% humidity in a dark room. After hatching in dark incubators, chicks were isolated and housed individually in metal cages (28 cm wide × 32 cm high × 40 cm deep) with food and water available *ad libitum*, at a constant room temperature of 30–32 °C and a constant light–dark regime of 14 h light and 10 h dark. All experimental protocols were approved by the research ethics committee of the University of Trento and by the Italian Ministry of Health (permit number 745/2021-PR).

### Experimental setup

The setup consisted of a rectangular shaped arena (28 X 40 X 32 cm; W X L X H) with metal walls that were grounded. In the centre of one of the shorter walls there was a circular opening (diameter 12 cm). A computer screen (AOC AGON AG271QG4, 144Hz) used for stimuli presentation was positioned directly behind the circular opening. Within the rectangular arena was a small wooden box 14 X 13 X 22 (W X L X H) cm whose frontal wall was made of a metal grid in. The box was placed 25 cm in front of the circular opening with the screen (Fig. 1A). During the experiments chicks remained inside the box from where they could observe the stimuli. Stimulus presentation was controlled by the PsychoPy toolbox (Pierce et al. 2019).

### Habituation procedure

The habituation occurred between the 3^rd^ and the 6^th^ day after hatching. On the 3^rd^ day post hatching, chicks learned to peck on mealworms. During the day 4 after hatching, chicks were, first, habituated to the experimental setup and then to the number stimuli appearing on the screen. The birds received mealworms every time after the stimulus appeared on the screen, which motivated them to pay attention to the moment when any stimulus would appear. The stimuli were presented and rewarded randomly, so that chicks would not associate any particular numerosity with the reward. During the days 5 and 6 post hatching we gradually decreased the reward rate, so that birds would still pay attention to the screen even without getting a mealworm. This procedure allowed us to minimize rewarding during actual recording sessions.

### Surgery and recordings

On the 7^th^ day after hatching chicks were fully anaesthetized using Isoflurane inhalation (1.5 – 2.0% gas volume, Vetflurane, 1000mg/g, Virbac, Italy) and placed in the stereotaxic apparatus with a bar fixed at the beaks’ base and tilted 45° to ear bars. Local anaesthesia (Emla cream, 2.5% lidocaine + 2.5% prilocaine, AstraZeneka, S.p.A.) was applied to the ears and skull skin before and after the surgery. Metal screws were placed into the skull for grounding and stabilisation of the implant. A small craniotomy was made in the skull on the right hemisphere above the NCL (1.0 mm anterior to the bregma, 4.5 mm lateral to the midline). For extracellular recordings we used self-wired tetrodes made out of formvar-insulated Nichrome wires (17.78 µm diameter, A-M Systems, USA), which were gold-plated to reduce the impedance to 300 – 400 kOm (controlled by nanoZ, Plexon Inc., USA). Then, a commercially available Halo-5 microdrive (Neuralynx, USA) was assembled according to the producer instructions, where four single tetrodes were put into polymicro tubes (inner diameter 0.1 mm) and glued to the plastic shuttles. The microdrive was implanted and fixed first with quick adhesive silicone (Kwik-Sil, World Precision Instruments, USA) and then with dental cement (Henry Schein Krugg Srl, Italy). To increase the probability of finding number-responsive units, we did not glue the electrode tips within the tetrodes We, thus, considered each tetrode as a brush-like arrangement of four single electrodes. Since in this brush arrangement the positions of the electrode tips can vary, some of the electrodes may have recorded signals from the same neurons. Hence, for sorting and subsequent analyses we chose only the best electrode from each tetrode at every recording position to avoid double-counting of the same neurons.

After the surgery, the chicks were left to recover until the next day in their home cages. Between the 8^th^ and the 12^th^ day after hatching we recorded neural responses to numerical stimuli in the NCL of chicks. Before every recording session the microdrive was connected to the Plexon system (Plexon Inc., USA) via a QuickClip connector and an omnetics headstage (Neuralynx, USA). After every recording session the tetrodes were manually advanced by 60 – 100 µm.

Signals were pre-amplified with a 16-channel head-stage (20×, Plexon Model number: PX.HST/16V-G20-LN) subsequently amplified 1000 ×, digitalised and filtered (300 Hz high-pass filter, 3 kHz low-pass filter and 50 Hz noise removal). Common average referencing (CAR) method (the averaged signal across channels) of the PlexControl system was used for referencing. Spikes were detected with the PlexControl software with an automatic 4-sigma threshold from the average noise level. Subsequently, spike sorting was performed manually in the Plexon Offline Sorter (see Figure S1 in supplementary information for examples of the raw signal and spike sorting).

### Stimuli

As numerical stimuli we used red dots outlined with a thin black line that appeared in the centre of the screen in a white background circle 6 cm in diameter. The size of stimuli ranged from 0.25 cm to 1.4 cm (0.6 – 3.2°, Schmid and Wildsoet 1998). We explored neural responses to numerosities from 1 to 5. To control for visual parameters that might interfere with numerosity perception, during every recording session we presented three different types of stimuli (Fig. 1B). “Radius-fixed” type of stimuli consisted of dots with a fixed radius, meaning that area and perimeter increase with numerosity. “Area-fixed” stimuli have constant total area over all numerosities, while the total perimeter of the dots increases with numerosity. “Perimeter-fixed” stimuli have constant total perimeter over all numerosities, while total area of these stimuli decreases with numerosity. To further control that neurons do not respond to other visual parameters except for quantity, the inter-distance interval between dots varied randomly. Moreover, for every day of recording we created a new batch of stimuli consisting of 30 unique images for each numerosity/stimulus type combination. Numerosity stimuli were created using GeNEsIS software (Zanon et al. 2021).

During recording sessions we randomly presented stimuli for 500 ms with 2000 ms of inter-stimulus interval. Experiments were video-recorded using CineLAB system (Plexon Inc., USA). To enhance the motivation of birds to pay attention to the screen, random trials were occasionally rewarded. These trials were subsequently discarded from the analysis.

### Histological analysis

After the last neural recording birds were overdosed with the ketamine/xylazine solution (1:1 ketamine 10 mg/ml + xylazine 2 mg/ml). Electrolytic lesions were made at the recording sites by applying a high-voltage current to the tetrodes for 10-15 seconds. Then, the birds were perfused intracardially with the phosphate buffer (PBS; 0.1 mol, pH = 7.4, 0.9% sodium chloride, 5 °C) followed by 4% paraformaldehyde (PFA). Brains were incubated for at least two days in PFA and further two days in 30% sucrose solution in PFA. Coronal 60 μm brain sections were cut at -20 °C using a cryostat (Leica CM1850 UV), mounted on glass slides, stained with the Giemsa dye (MG500, Sigma-Aldrich, St. Louis, USA), and cover slipped with Eukitt (FLUKA). Brain sections were examined under the stereomicroscope (Stemi 508, Carl Zeiss, Oberkochen, Germany) to estimate the anatomical position of recording sites.

### Data analysis

Based on the analysis of video-recordings, we selected only those trials, when birds were not rewarded. Since in birds there is an almost complete decussation of the optic fibres we recorded from the right hemisphere and we selected only trials, when birds looked at the stimulus with both eyes or with the contralateral (left) eye. For each recorded unit, we excluded trials with a firing rate of less than 1 Hz. For the final analysis we considered only those units that were recorded for at least 7 trials for each numerosity and stimulus type (on average 23 trials per numerosity/stimulus type).

The neural activity of recorded units was analysed as the mean firing rate over 500 ms of stimulus presentation. To find numerosity-responsive neurons we performed two-way ANOVA with the numerosity (1 to 5) and the stimulus type (“radius-fixed”, “area-fixed”, “perimeter-fixed”) as factors. We considered the recorded neuron as numerosity-responsive if only the effect of the “numerosity” factor was highly significant (p<0.01), but not of the stimulus type or the interaction between the two factors. The numerosity that elicited the strongest neural response was defined as a preferred numerosity for this neuron.

For every number neuron we performed a post hoc analysis (Tukey’s test) to compare the neural response between the most preferred and other numerosities. To test for the numerical distance effect we applied a GLM for binomial data with the logit-link function. In the model, the proportion of neurons that significantly differentiate between given numerosities was taken as the response variable, and either the absolute numerical distance or the logarithm of the numerical distance was taken as a factor. We then compared the goodness-of-fit of these two models based on the difference in the Bayesian Information Criterion (ΔBIC), where the lower is the BIC value, the better is the model’s fit.

To validate the stability of our recordings we performed a cross-validation analysis. For each numerosity-responsive unit we calculated the preferred numerosity for the first and the second half of all trials separately. If the neural response to number stimuli was stable across the recording, we expected the Pearson’s correlation between the first and the second half of the trials to be close to 1 for the whole population of number neurons. The cross-validation analysis showed a strong correlation of 0.82 (p<0.001) between the preferred numerosity in the first and the second half of the trials, confirming the stability of our recordings.

For every number neuron we normalised neural activity by setting the firing rate in response to the preferred numerosity at 100% and to the least preferred numerosity at 0%. The resulting neural filter functions were averaged by group based on the preferred numerosity, thus, creating numerosity tuning curves for e.g. neurons preferring numerosity 1, numerosity 2, etc.

To evaluate the chance level of finding false-positive numerical responses in our dataset, for every recorded neuron we shuffled all the trials for 1000 times and performed an ANOVA each time to select false-positive number neurons. We compared the proportion of false-positive and real number neurons with the proportion test. We further compared the tuning curves between the false-positive number neurons and the real number neurons. For this, we randomly sampled from the false-positive neurons the same number of neurons as the one of actually recorded number neurons. We compared the tuning curves of real and false-positive neurons performing a two-factor ANOVA with the interaction of “numerical distance” and “data type” (real/false positive).

According to the Weber-Fechner law, the perception of sensory stimuli (including quantities) is proportional to a logarithmic (Fechner 1860) or power scale (Stevens 1961) of stimulus magnitude. Therefore, when plotted on a linear scale, one might expect tuning curves to become increasingly asymmetric and wide with increasing numerosity. These properties are usually referred to as a numerical distance effect and a numerical magnitude effect respectively. To evaluate the symmetry and the width of the neural filter functions we fitted the Gaussian function to the curves (MatLab Curve Fitting Toolbox) plotted on four different scales: linear, a power function with an exponent of 0.5, a power function with an exponent of 0.33, and a logarithmic (log2) scale (Ditz and Nieder 2016). The symmetry of the Gaussian fit was estimated based on R-squared (*r^2^*) values, i.e. the higher is *r^2^* the better and more symmetric is the fit. The width of the Gaussian fit was reflected by its sigma (σ). The four scaling methods were compared based on the *r^2^* values by the Friedman’s test for non-parametric data with repeated measures with the post-hoc pairwise comparison with the Nemenyi test. The relationship between the numerosity and the sigma of the Gaussian was tested by an ANOVA for different scaling methods separately.

All statistical analyses and visualization of the data was performed in R (R Core Team 2020) with packages “tidyverse”, “ggplot2”, and “PMCMRplus” and in MATLAB using custom-made scripts and the Curve Fitting Toolbox.

## Supporting information

Supplementary Data Set

## Acknowledgments and funding sources

We are grateful to Anastasia Morandi-Raikova for her help with handling the chicks. Elena Lorenzi, Andrea Messina, and Matilde Perrino provided very valuable comments to the current manuscript. This work was supported by funding from the European Research Council (ERC) under the European Union’s Horizon 2020 research and innovation program (Grant agreement no. 833504 SPANUMBRA) and from a PRIN 2017 ERC-SH4–A (2017PSRHPZ) to G. V.

## Author contributions

D.K, U.M. and G.V. designed the experiment. D.K. and U.M. established the procedure for behavioural training and electrophysiology in freely moving chicks. D.K. performed the experiment and analysed the data. M.Z. developed the numerical stimuli and contributed to apart of the data analysis. D.K. wrote the original draft of the manuscript and implemented the comments of all authors. U.M. and G.V. revised and edited the manuscript.

## Availability of data and material

A summary table including the mean firing rates for all numerosity-responsive neurons for all trials is included as a supplementary data file. Any additional data that support the findings of this study are available from the corresponding author upon reasonable request.

## Code availability

The custom codes that were used for data analysis are available from the corresponding author upon reasonable request.

## Supplementary information

**Figure S1.**
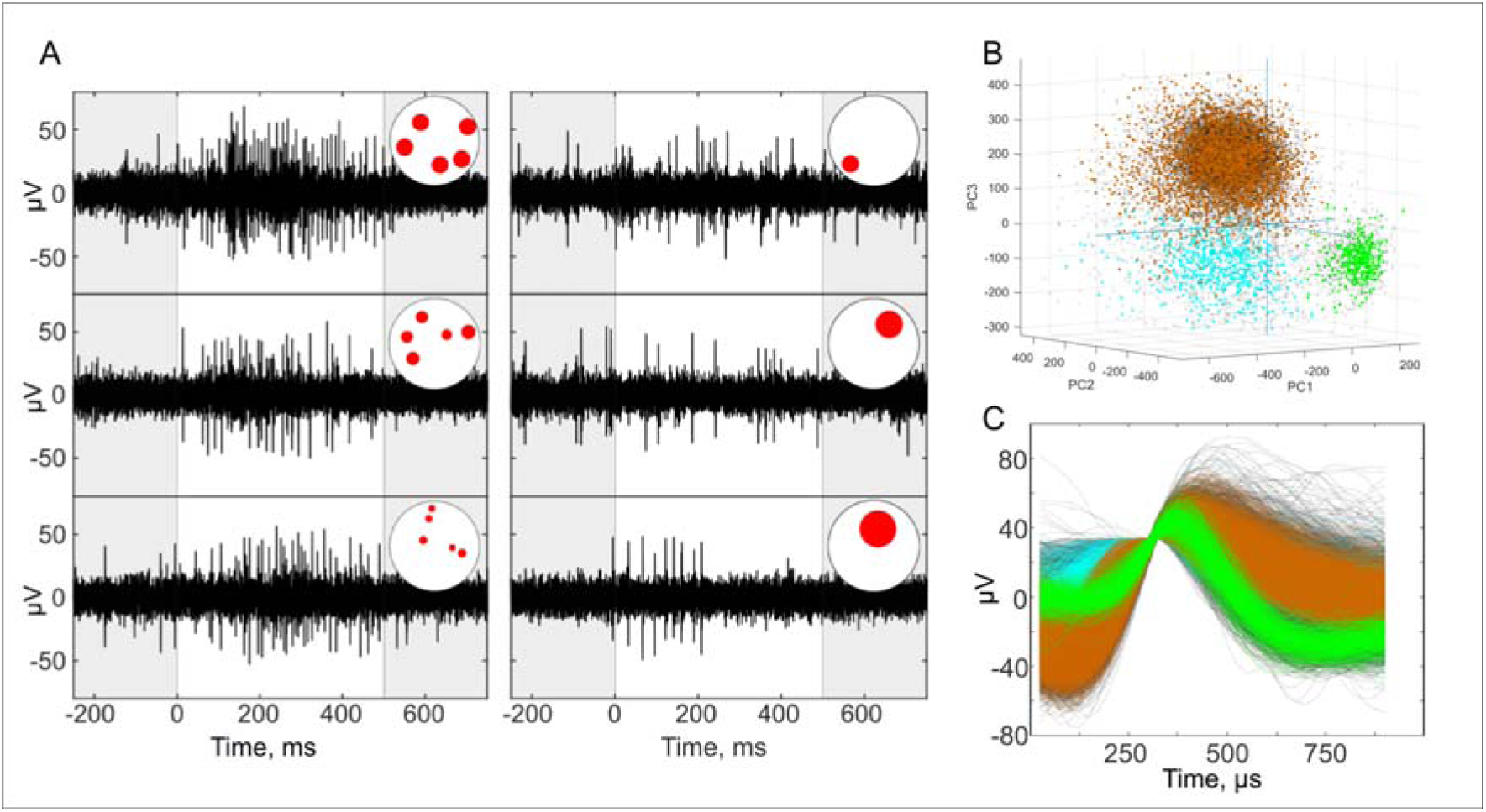
(A) Example of raw spike trains: electrical signal is shown after the high-pass filter was applied. Examples of single trials, representing neural response of the unit to numerosity 5 (left column), irrespective of the stimulus appearance (top: radius-fixed, middle: area-fixed, bottom: perimeter-fixed). Note the decreasing neural response to the numerosity 1 (right column). (B) The PCA clustering of the corresponding recording with waveforms of different units shown by different colours. The waveforms of the number-responsive unit are shown in orange, unsorted waveforms are shown in grey. (C) Spike waveforms of corresponding units isolated from the recording of one electrode.

**Table S1.** Summary of the two-way ANOVA for every recorded unit. Unit: id of the recorded unit. Preferred numerosity: numerosity stimulus that elicited the strongest response in the corresponding unit. ANOVA results (F-statistics and p-value) are summarized for the factor “Stimulus type” (radius-fixed, area-fixed, perimeter-fixed), “Numerosity” (numerosity “one” to “five”), or interaction between them.

| Unit | Preferred numerosity | ANOVA (Firing rate ~ Stimulus type * Numerosity) |  |  |
| --- | --- | --- | --- | --- |
|  |  | Numerosity | Stimulus type | Interaction |
| 1 | num1 | F(4,546)=4.0327, p = 0.003 | F(2,546)=0.5228, p = 0.593 | F(8,546)=1.4242, p = 0.183 |
| 2 | num1 | F(4,197)=3.5672, p = 0.008 | F(2,197)=0.4656, p = 0.628 | F(8,197)=0.8763, p = 0.537 |
| 3 | num1 | F(4,202)=4.0268, p = 0.004 | F(2,202)=1.7732, p = 0.172 | F(8,202)=1.1967, p = 0.303 |
| 4 | num1 | F(4,565)=4.2588, p = 0.002 | F(2,565)=0.0012, p = 0.999 | F(8,565)=0.7852, p = 0.616 |
| 5 | num1 | F(4,270)=5.3565, p = 0 | F(2,270)=1.2038, p = 0.302 | F(8,270)=0.9519, p = 0.474 |
| 6 | num1 | F(4,270)=5.4507, p = 0 | F(2,270)=0.0475, p = 0.954 | F(8,270)=0.7056, p = 0.687 |
| 7 | num1 | F(4,239)=3.7861, p = 0.005 | F(2,239)=0.4114, p = 0.663 | F(8,239)=1.6597, p = 0.109 |
| 8 | num1 | F(4,419)=4.9175, p = 0.001 | F(2,419)=1.0111, p = 0.365 | F(8,419)=1.0988, p = 0.363 |
| 9 | num1 | F(4,399)=7.4226, p = 0 | F(2,399)=2.8675, p = 0.058 | F(8,399)=2.1769, p = 0.028 |
| 10 | num1 | F(4,366)=4.2597, p = 0.002 | F(2,366)=0.9588, p = 0.384 | F(8,366)=1.4233, p = 0.185 |
| 11 | num1 | F(4,364)=6.4075, p = 0 | F(2,364)=0.2249, p = 0.799 | F(8,364)=1.2606, p = 0.263 |
| 12 | num1 | F(4,490)=3.9053, p = 0.004 | F(2,490)=1.1902, p = 0.305 | F(8,490)=0.6286, p = 0.754 |
| 13 | num1 | F(4,251)=5.7273, p = 0 | F(2,251)=0.9275, p = 0.396 | F(8,251)=1.2411, p = 0.273 |
| 14 | num1 | F(4,186)=3.5545, p = 0.008 | F(2,186)=1.1998, p = 0.304 | F(8,186)=0.6334, p = 0.749 |
| 15 | num1 | F(4,233)=4.4063, p = 0.002 | F(2,233)=1.138, p = 0.322 | F(8,233)=0.3761, p = 0.933 |
| 16 | num1 | F(4,316)=3.6533, p = 0.006 | F(2,316)=1.4135, p = 0.245 | F(8,316)=0.8235, p = 0.582 |
| 17 | num2 | F(4,177)=4.5348, p = 0.002 | F(2,177)=0.2742, p = 0.761 | F(8,177)=0.6772, p = 0.711 |
| 18 | num2 | F(4,176)=4.1623, p = 0.003 | F(2,176)=0.8853, p = 0.414 | F(8,176)=0.8814, p = 0.533 |
| 19 | num2 | F(4,147)=3.6633, p = 0.007 | F(2,147)=0.929, p = 0.397 | F(8,147)=0.5183, p = 0.841 |
| 20 | num2 | F(4,145)=3.7461, p = 0.006 | F(2,145)=0.514, p = 0.599 | F(8,145)=0.5375, p = 0.827 |
| 21 | num2 | F(4,298)=3.4934, p = 0.008 | F(2,298)=0.7184, p = 0.488 | F(8,298)=1.0979, p = 0.364 |
| 22 | num2 | F(4,253)=3.5881, p = 0.007 | F(2,253)=0.2355, p = 0.79 | F(8,253)=0.454, p = 0.887 |
| 23 | num3 | F(4,213)=3.4856, p = 0.009 | F(2,213)=0.6473, p = 0.524 | F(8,213)=0.5014, p = 0.854 |
| 24 | num3 | F(4,184)=3.6183, p = 0.007 | F(2,184)=0.5629, p = 0.571 | F(8,184)=0.6608, p = 0.725 |
| 25 | num3 | F(4,472)=5.1242, p = 0 | F(2,472)=2.8061, p = 0.062 | F(8,472)=0.809, p = 0.595 |
| 26 | num3 | F(4,410)=3.7563, p = 0.005 | F(2,410)=4.5228, p = 0.011 | F(8,410)=1.3146, p = 0.233 |
| 27 | num3 | F(4,496)=5.8789, p = 0 | F(2,496)=0.1179, p = 0.889 | F(8,496)=1.4538, p = 0.173 |
| 28 | num4 | F(4,176)=3.926, p = 0.004 | F(2,176)=1.1196, p = 0.329 | F(8,176)=0.6298, p = 0.752 |
| 29 | num4 | F(4,178)=3.9175, p = 0.004 | F(2,178)=0.4244, p = 0.655 | F(8,178)=0.6897, p = 0.7 |
| 30 | num4 | F(4,267)=4.5852, p = 0.001 | F(2,267)=0.6043, p = 0.547 | F(8,267)=0.8609, p = 0.55 |
| 31 | num4 | F(4,371)=3.6503, p = 0.006 | F(2,371)=1.9928, p = 0.138 | F(8,371)=0.5894, p = 0.787 |
| 32 | num4 | F(4,430)=3.4615, p = 0.008 | F(2,430)=0.3739, p = 0.688 | F(8,430)=0.513, p = 0.847 |
| 33 | num4 | F(4,420)=4.4972, p = 0.001 | F(2,420)=1.1432, p = 0.32 | F(8,420)=1.1135, p = 0.353 |
| 34 | num4 | F(4,459)=6.5488, p = 0 | F(2,459)=2.8234, p = 0.06 | F(8,459)=1.3186, p = 0.232 |
| 35 | num4 | F(4,333)=3.4399, p = 0.009 | F(2,333)=3.3226, p = 0.037 | F(8,333)=1.5331, p = 0.145 |
| 36 | num4 | F(4,308)=4.2385, p = 0.002 | F(2,308)=0.1141, p = 0.892 | F(8,308)=0.9124, p = 0.506 |
| 37 | num5 | F(4,323)=5.0184, p = 0.001 | F(2,323)=1.9782, p = 0.14 | F(8,323)=0.3722, p = 0.935 |
| 38 | num5 | F(4,188)=3.5707, p = 0.008 | F(2,188)=1.1655, p = 0.314 | F(8,188)=0.5456, p = 0.821 |
| 39 | num5 | F(4,307)=5.1489, p = 0 | F(2,307)=0.0322, p = 0.968 | F(8,307)=0.4935, p = 0.861 |
| 40 | num5 | F(4,260)=5.1953, p = 0 | F(2,260)=0.945, p = 0.39 | F(8,260)=0.682, p = 0.707 |
| 41 | num5 | F(4,234)=4.65, p = 0.001 | F(2,234)=0.7971, p = 0.452 | F(8,234)=0.8709, p = 0.542 |
| 42 | num5 | F(4,232)=3.7216, p = 0.006 | F(2,232)=0.0802, p = 0.923 | F(8,232)=1.3969, p = 0.199 |
| 43 | num5 | F(4,484)=3.6509, p = 0.006 | F(2,484)=0.1205, p = 0.887 | F(8,484)=0.7686, p = 0.631 |
| 44 | num5 | F(4,458)=12.8202, p = 0 | F(2,458)=0.7574, p = 0.469 | F(8,458)=0.4219, p = 0.908 |
| 45 | num5 | F(4,192)=4.4432, p = 0.002 | F(2,192)=2.0198, p = 0.135 | F(8,192)=1.8229, p = 0.075 |
| 46 | num5 | F(4,415)=3.8044, p = 0.005 | F(2,415)=1.618, p = 0.2 | F(8,415)=0.5316, p = 0.832 |
| 47 | num5 | F(4,685)=14.1207, p = 0 | F(2,685)=1.9687, p = 0.141 | F(8,685)=1.4958, p = 0.156 |
| 48 | num5 | F(4,371)=6.7422, p = 0 | F(2,371)=0.6683, p = 0.513 | F(8,371)=1.6257, p = 0.115 |
| 49 | num5 | F(4,359)=4.1619, p = 0.003 | F(2,359)=0.1355, p = 0.873 | F(8,359)=0.4252, p = 0.906 |
| 50 | num5 | F(4,355)=13.4938, p = 0 | F(2,355)=1.832, p = 0.162 | F(8,355)=0.3219, p = 0.958 |
| 51 | num5 | F(4,354)=6.4972, p = 0 | F(2,354)=0.6433, p = 0.526 | F(8,354)=0.9821, p = 0.45 |
| 52 | num5 | F(4,288)=3.3897, p = 0.01 | F(2,288)=2.1069, p = 0.123 | F(8,288)=1.2445, p = 0.273 |
| 53 | num5 | F(4,460)=5.4865, p = 0 | F(2,460)=1.4027, p = 0.247 | F(8,460)=0.7622, p = 0.636 |

**Table S2.**
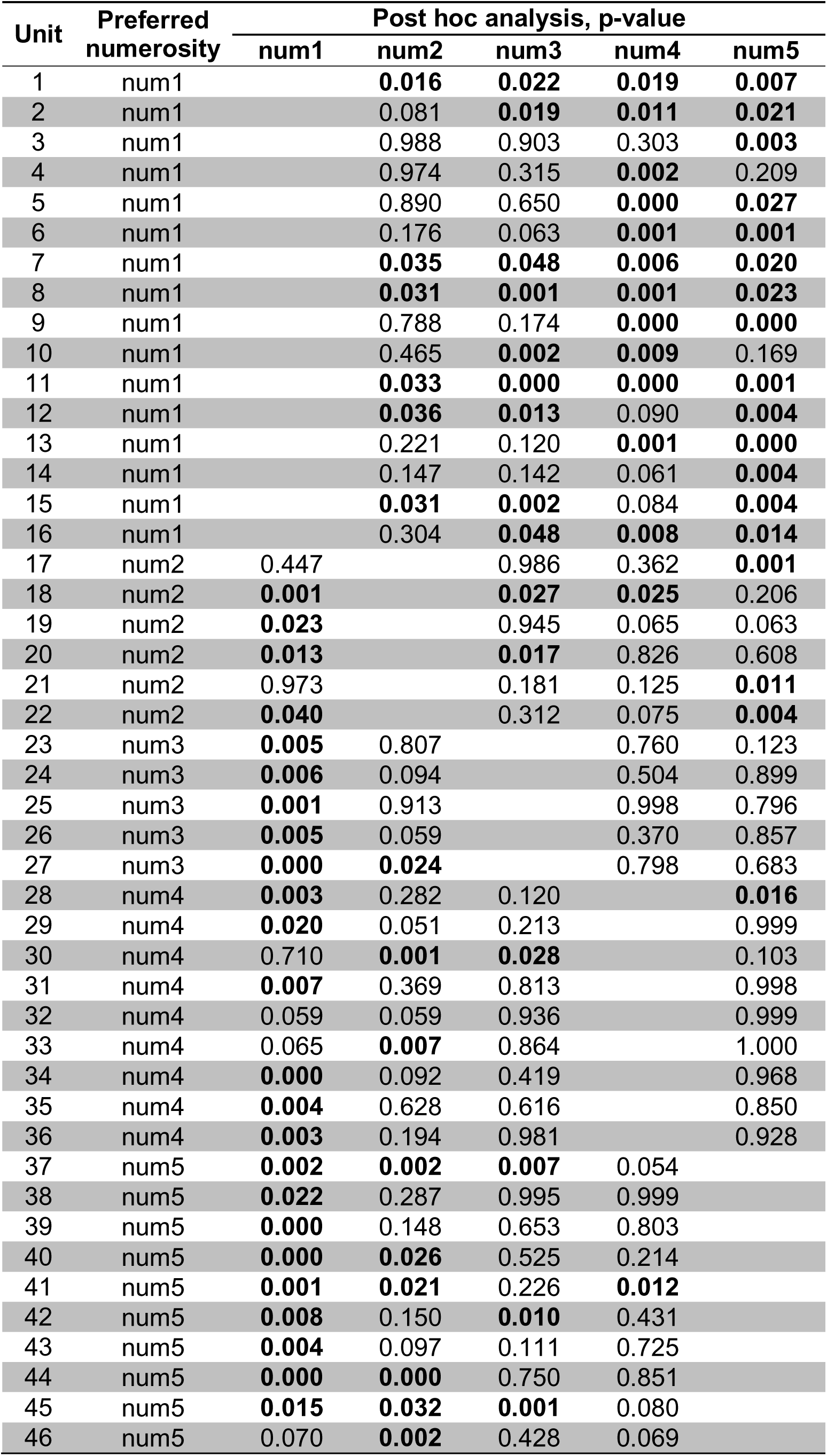
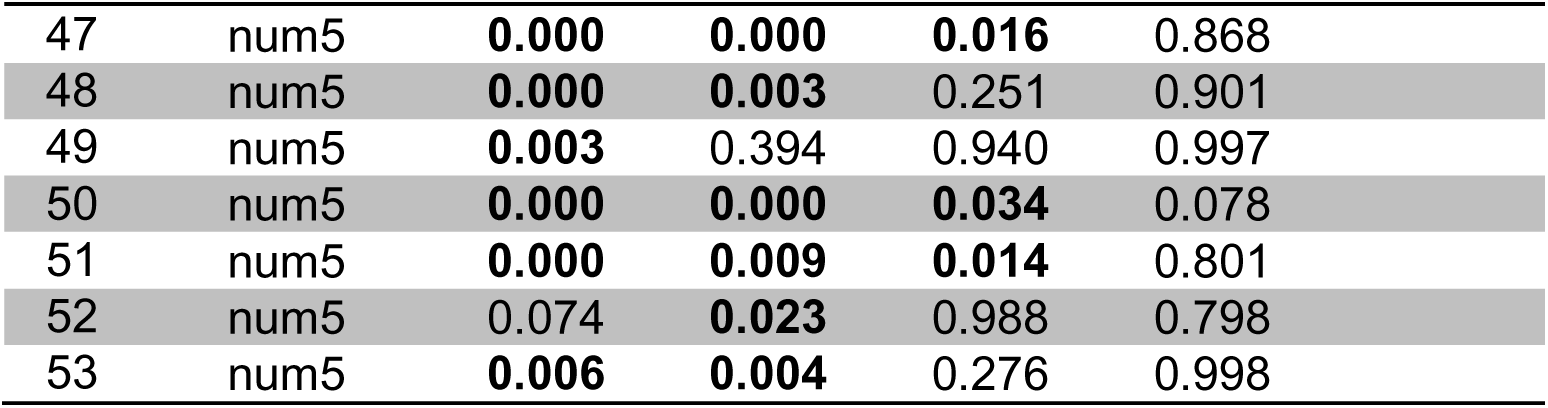
The post hoc analysis of the two-way ANOVA based on the Tukey-Kramer method summarising p-values for every pairwise comparison between the most-preferred and other numerosities. Significant p-values <0.05 are highlighted in bold.

| Unit | Preferred numerosity | Post hoc analysis, p-value |  |  |  |  |
| --- | --- | --- | --- | --- | --- | --- |
|  |  | num1 | num2 | num3 | num4 | num5 |
| 1 | num1 |  | **0.016** | **0.022** | **0.019** | **0.007** |
| 2 | num1 |  | 0.081 | **0.019** | **0.011** | **0.021** |
| 3 | num1 |  | 0.988 | 0.903 | 0.303 | **0.003** |
| 4 | num1 |  | 0.974 | 0.315 | **0.002** | 0.209 |
| 5 | num1 |  | 0.890 | 0.650 | **0.000** | **0.027** |
| 6 | num1 |  | 0.176 | 0.063 | **0.001** | **0.001** |
| 7 | num1 |  | **0.035** | **0.048** | **0.006** | **0.020** |
| 8 | num1 |  | **0.031** | **0.001** | **0.001** | **0.023** |
| 9 | num1 |  | 0.788 | 0.174 | **0.000** | **0.000** |
| 10 | num1 |  | 0.465 | **0.002** | **0.009** | 0.169 |
| 11 | num1 |  | **0.033** | **0.000** | **0.000** | **0.001** |
| 12 | num1 |  | **0.036** | **0.013** | 0.090 | **0.004** |
| 13 | num1 |  | 0.221 | 0.120 | **0.001** | **0.000** |
| 14 | num1 |  | 0.147 | 0.142 | 0.061 | **0.004** |
| 15 | num1 |  | **0.031** | **0.002** | 0.084 | **0.004** |
| 16 | num1 |  | 0.304 | **0.048** | **0.008** | **0.014** |
| 17 | num2 | 0.447 |  | 0.986 | 0.362 | **0.001** |
| 18 | num2 | **0.001** |  | **0.027** | **0.025** | 0.206 |
| 19 | num2 | **0.023** |  | 0.945 | 0.065 | 0.063 |
| 20 | num2 | **0.013** |  | **0.017** | 0.826 | 0.608 |
| 21 | num2 | 0.973 |  | 0.181 | 0.125 | **0.011** |
| 22 | num2 | **0.040** |  | 0.312 | 0.075 | **0.004** |
| 23 | num3 | **0.005** | 0.807 |  | 0.760 | 0.123 |
| 24 | num3 | **0.006** | 0.094 |  | 0.504 | 0.899 |
| 25 | num3 | **0.001** | 0.913 |  | 0.998 | 0.796 |
| 26 | num3 | **0.005** | 0.059 |  | 0.370 | 0.857 |
| 27 | num3 | **0.000** | **0.024** |  | 0.798 | 0.683 |
| 28 | num4 | **0.003** | 0.282 | 0.120 |  | **0.016** |
| 29 | num4 | **0.020** | 0.051 | 0.213 |  | 0.999 |
| 30 | num4 | 0.710 | **0.001** | **0.028** |  | 0.103 |
| 31 | num4 | **0.007** | 0.369 | 0.813 |  | 0.998 |
| 32 | num4 | 0.059 | 0.059 | 0.936 |  | 0.999 |
| 33 | num4 | 0.065 | **0.007** | 0.864 |  | 1.000 |
| 34 | num4 | **0.000** | 0.092 | 0.419 |  | 0.968 |
| 35 | num4 | **0.004** | 0.628 | 0.616 |  | 0.850 |
| 36 | num4 | **0.003** | 0.194 | 0.981 |  | 0.928 |
| 37 | num5 | **0.002** | **0.002** | **0.007** | 0.054 |  |
| 38 | num5 | **0.022** | 0.287 | 0.995 | 0.999 |  |
| 39 | num5 | **0.000** | 0.148 | 0.653 | 0.803 |  |
| 40 | num5 | **0.000** | **0.026** | 0.525 | 0.214 |  |
| 41 | num5 | **0.001** | **0.021** | 0.226 | **0.012** |  |
| 42 | num5 | **0.008** | 0.150 | **0.010** | 0.431 |  |
| 43 | num5 | **0.004** | 0.097 | 0.111 | 0.725 |  |
| 44 | num5 | **0.000** | **0.000** | 0.750 | 0.851 |  |
| 45 | num5 | **0.015** | **0.032** | **0.001** | 0.080 |  |
| 46 | num5 | 0.070 | **0.002** | 0.428 | 0.069 |  |

|  |  |  |  |  |  |
| --- | --- | --- | --- | --- | --- |
| 47 | num5 | **0.000** | **0.000** | **0.016** | 0.868 |
| 48 | num5 | **0.000** | **0.003** | 0.251 | 0.901 |
| 49 | num5 | **0.003** | 0.394 | 0.940 | 0.997 |
| 50 | num5 | **0.000** | **0.000** | **0.034** | 0.078 |
| 51 | num5 | **0.000** | **0.009** | **0.014** | 0.801 |
| 52 | num5 | 0.074 | **0.023** | 0.988 | 0.798 |
| 53 | num5 | **0.006** | **0.004** | 0.276 | 0.998 |

## References

Balestrieri, A., Gazzola, A., Pellitteri-Rosa, D. & Vallortigara, G. Discrimination of group numerousness under predation risk in anuran tadpoles. Anim. Cogn. 22(2), 223–230 (2019). https://doi.org/10.1007/s10071-019-01238-5

Beran, M. J., Evans, T. A. & Harris, E. H. Perception of food amounts by chimpanzees based on the number, size, contour length and visibility of items. Anim. Behav. 75, 1793–1802 (2008). https://doi.org/10.1016/j.anbehav.2007.10.035

Bortot, M., Regolin, L. & Vallortigara, G. A sense of number in invertebrates. Biochem. Biophys. Res. Commun. 564, 37–42 (2021). https://doi.org/10.1016/j.bbrc.2020.11.039

Brannon, E. M. & Merritt, D. J. Evolutionary foundations of the approximate number system. in Space, time and number in the brain (eds. Dehaene, S. & Brannon, E.M.) 207–224 (Academic Press, 2011). https://doi.org/10.1016/B978-0-12-385948-8.00014-1

Briscoe, S. D. & Ragsdale, C. W. Evolution of the chordate telencephalon. Curr. Biol. 29(13), 647–662 (2019). https://doi.org/10.1016/j.cub.2019.05.026

Collins, E., Park, J. & Behrmann, M. Numerosity representation is encoded in human subcortex. Proc. Natl Acad. Sci. USA 114(14), 2806–2815 (2017). https://doi.org/10.1073/pnas.1613982114

Davis, H. & Albert, M. Numerical discrimination by rats using sequential auditory stimuli. Animal Learning & Behavior 14(1), 57–59 (1986). https://doi.org/10.3758/BF03200037

Diekamp, B., Kalt, T. & Güntürkün, O. Working memory neurons in pigeons. J. Neurosci. 22(4), RC210-R210 (2002). https://doi.org/10.1523/JNEUROSCI.22-04-j0002.2002

Ditz, H. M. & Nieder, A. Neurons selective to the number of visual items in the corvid songbird endbrain. Proc. Natl Acad. Sci. USA 112(25), 7827–7832 (2015). https://doi.org/10.1073/pnas.1504245112

Ditz, H. M. & Nieder, A. Numerosity representations in crows obey the Weber–Fechner law. Proc. R. Soc. B Biol. Sci. 283(1827), 20160083 (2016). https://doi.org/10.1098/rspb.2016.0083

Fechner, G. T. Elemente der psychophysik (Vol. 2). (Breitkopf u. Härtel, 1860).

Gazzola, A., Vallortigara, G. & Pellitteri-Rosa, D. Continuous and discrete quantity discrimination in tortoises. Biol. Lett. 14(12), 20180649 (2018). https://doi.org/10.1098/rsbl.2018.0649

Güntürkün, O. & Bugnyar, T. Cognition without cortex. *Trends Cogn. Sci.* **20**(4), 291–303 (2016). https://doi.org/10.1016/j.tics.2016.02.001

Güntürkün, O., von Eugen, K., Packheiser, J. & Pusch, R. Avian pallial circuits and cognition: A comparison to mammals. Curr. Opin. Neurobiol. 71, 29–36 (2021). https://doi.org/10.1016/j.conb.2021.08.007

Gusel’nikov, V. I., Morenkov, É. D. & Gutsu, I. P. Responses of neurons in the pigeon’s optic tectum to visual stimuli. Neurophysiology 3(1), 78–83 (1971).

Hahn, L. A., Balakhonov, D., Fongaro, E., Nieder, A. & Rose, J. Working memory capacity of crows and monkeys arises from similar neuronal computations. eLife 10, e72783 (2021). DOI: 10.7554/eLife.72783

Hunt, S., Low, J. & Burns, K. C. Adaptive numerical competency in a food-hoarding songbird. Proc. R. Soc. B Biol. Sci. 275(1649), 2373–2379 (2008). https://doi.org/10.1098/rspb.2008.0702

Hussar, C. R. & Pasternak, T. Flexibility of sensory representations in prefrontal cortex depends on cell type. Neuron 64(5), 730–743 (2009). https://doi.org/10.1016/j.neuron.2009.11.018

Izard, V., Sann, C., Spelke, E. S. & Streri, A. Newborn infants perceive abstract numbers. Proc. Natl. Acad. Sci. USA 106(25), 10382–10385 (2009). https://doi.org/10.1073/pnas.0812142106

Kim, G., Jang, J., Baek, S., Song, M. & Paik, S. B. Visual number sense in untrained deep neural networks. Sci. Adv. 7(1), eabd6127 (2021). DOI: 10.1126/sciadv.abd6127

Kutter, E. F., Bostroem, J., Elger, C. E., Mormann, F. & Nieder, A. Single neurons in the human brain encode numbers. Neuron 100(3), 753–761 (2018). https://doi.org/10.1016/j.neuron.2018.08.036

Lorenzi, E., Perrino, M. & Vallortigara, G. Numerosities and Other Magnitudes in the Brains: A Comparative View. Front. Psychol. 12, 1104 (2021). https://doi.org/10.3389/fpsyg.2021.641994

Lyon, B. E. Egg recognition and counting reduce costs of avian conspecific brood parasitism. Nature 422, 495–499 (2003). https://doi.org/10.1038/nature01505

Messina, A., Potrich, D., Schiona, I., Sovrano, V.A., Fraser, S.E., Brennan, C.H. & Vallortigara, G. Neurons in the dorso-central division of zebrafish pallium respond to change in visual numerosity. *Cereb. Cortex*., bhab218 (2021). DOI: 10.1093/cercor/bhab218

Nasr, K., Viswanathan, P. & Nieder, A. Number detectors spontaneously emerge in a deep neural network designed for visual object recognition. Sci. Adv. 5(5), eaav7903 (2019). DOI: 10.1126/sciadv.aav7903

Nieder, A. Supramodal numerosity selectivity of neurons in primate prefrontal and posterior parietal cortices. Proc. Natl. Acad. Sci. USA 109(29), 11860–11865 (2012). https://doi.org/10.1073/pnas.1204580109

Nieder, A. The neuronal code for number. Nat. Rev. Neurosci. 17(6), 366–382 (2016). https://doi.org/10.1038/nrn.2016.40

Nieder, A. A brain for numbers: the biology of the number instinct. (MIT press, 2019).

Nieder, A., Freedman, D. J. & Miller, E. K. Representation of the quantity of visual items in the primate prefrontal cortex. Science 297(5587), 1708–1711 (2002). DOI: 10.1126/science.1072493

Nieder, A., Wagener, L. & Rinnert, P. A neural correlate of sensory consciousness in a corvid bird. Science 369(6511), 1626–1629 (2020). DOI: 10.1126/science.abb1447

Olkowicz, S., Kocourek, M., Lučan, R.K., Porteš, M., Fitch, W.T., Herculano-Houzel, S. & Němec, P. Birds have primate-like numbers of neurons in the forebrain. Proc. Natl. Acad. Sci. USA 113(26), 7255–7260 (2016). https://doi.org/10.1073/pnas.1517131113

Peirce, J., Gray, J.R., Simpson, S., MacAskill, M., Höchenberger, R., Sogo, H., Kastman, E. and Lindeløv, J.K. PsychoPy2: Experiments in behavior made easy. Behav. Res. Methods 51(1), 195–203 (2019). https://doi.org/10.3758/s13428-018-01193-y

Potrich, D., Sovrano, V. A., Stancher, G. & Vallortigara, G. Quantity discrimination by zebrafish (*Danio rerio*). J. Comp. Psychol. 129(4), 388 (2015). https://doi.org/10.1037/com0000012

Preuss, S. J., Trivedi, C. A., vom Berg-Maurer, C. M., Ryu, S. & Bollmann, J. H. Classification of object size in retinotectal microcircuits. Curr. Biol. 24(20), 2376–2385 (2014). https://doi.org/10.1016/j.cub.2014.09.012

Preuss, T. M. & Wise, S. P. Evolution of prefrontal cortex. Neuropsychopharmacology 47(1), 3–19 (2022). https://doi.org/10.1038/s41386-021-01076-5

Prum, R.O., Berv, J.S., Dornburg, A., Field, D.J., Townsend, J.P., Lemmon, E.M. & Lemmon, A.R. A comprehensive phylogeny of birds (Aves) using targeted next-generation DNA sequencing. Nature 526(7574), 569-573 (2015). https://doi.org/10.1038/nature15697

Puelles, L. Comments on the updated tetrapartite pallium model in the mouse and chick, featuring a homologous claustro-insular complex. Brain Behav. Evol. 90(2), 171–189 (2017). https://doi.org/10.1159/000479782

R Core Team (2020). R: A language and environment for statistical computing. R Foundation for Statistical Computing, Vienna, Austria. URL https://www.R-project.org/

Rugani, R., Cavazzana, A., Vallortigara, G. & Regolin, L. One, two, three, four, or is there something more? Numerical discrimination in day-old domestic chicks. Anim. Cogn. 16(4), 557–564 (2013). DOI 10.1007/s10071-012-0593-8

Rugani, R., Fontanari, L., Simoni, E., Regolin, L. & Vallortigara, G. Arithmetic in newborn chicks. Proc. R. Soc. B Biol. Sci. 276(1666), 2451-2460 (2009). https://doi.org/10.1098/rspb.2009.0044

Rugani, R., Regolin, L. & Vallortigara, G. Discrimination of small numerosities in young chicks. J. Exp. Psychol. Anim. Behav. Process. 34(3), 388 (2008). https://doi.org/10.1037/0097-7403.34.3.388

Rugani, R., Vallortigara, G., Priftis, K. & Regolin, L. Number-space mapping in the newborn chick resembles humans’ mental number line. Science 347(6221), 534-536 (2015). DOI: 10.1126/science.aaa1379

Sawamura, H., Shima, K. & Tanji, J. Numerical representation for action in the parietal cortex of the monkey. Nature 415(6874), 918-922 (2002). https://doi.org/10.1038/415918a

Schmid, K. L. & Wildsoet, C. F. Assessment of visual acuity and contrast sensitivity in the chick using an optokinetic nystagmus paradigm. Vision Res. 38(17), 2629–2634 (1998). https://doi.org/10.1016/S0042-6989(97)00446-X

Stacho, M., Herold, C., Rook, N., Wagner, H., Axer, M., Amunts, K. & Güntürkün, O. A cortex-like canonical circuit in the avian forebrain. Science 369(6511), (2020). DOI: 10.1126/science.abc5534

Stancher, G., Rugani, R., Regolin, L. & Vallortigara, G. Numerical discrimination by frogs (*Bombina orientalis*). Anim. Cogn. 18(1), 219–229 (2015). DOI 10.1007/s10071-014-0791-7

Stevens, S. S. To honor Fechner and repeal his law. Science 133(3446), 80–86 (1961).

Striedter, G. F. & Northcutt, R. G. Brains through time: a natural history of vertebrates. (Oxford University Press, 2019).

Templeton, C. N., Greene, E. & Davis, K. Allometry of alarm calls: black-capped chickadees encode information about predator size. Science 308, 1934–1937 (2005). DOI: 10.1126/science.1108841

Veit, L. & Nieder, A. Abstract rule neurons in the endbrain support intelligent behaviour in corvid songbirds. Nat. Commun. 4(1), 1–11 (2013). https://doi.org/10.1038/ncomms3878

Veit, L., Pidpruzhnykova, G. & Nieder, A. Associative learning rapidly establishes neuronal representations of upcoming behavioral choices in crows. Proc. Natl. Acad. Sci. USA 112(49), 15208-15213 (2015). https://doi.org/10.1073/pnas.1509760112

Viswanathan, P. & Nieder, A. Neuronal correlates of a visual “sense of number” in primate parietal and prefrontal cortices. Proc. Natl. Acad. Sci. USA 110(27), 11187–11192 (2013). https://doi.org/10.1073/pnas.1308141110

von Eugen, K., Tabrik, S., Güntürkün, O. & Ströckens, F. A comparative analysis of the dopaminergic innervation of the executive caudal nidopallium in pigeon, chicken, zebra finch, and carrion crow. J. Comp. Neurol. 528(17), 2929–2955 (2020). https://doi.org/10.1002/cne.24878

Wagener, L., Loconsole, M., Ditz, H. M. & Nieder, A. Neurons in the endbrain of numerically naive crows spontaneously encode visual numerosity. Curr. Biol. 28(7), 1090–1094 (2018). https://doi.org/10.1016/j.cub.2018.02.023

Ward, C. & Smuts, B. Quantity-based judgments in the domestic dog (*Canis lupus familiaris*). Anim. Cogn. 10, 71–80 (2007). DOI 10.1007/s10071-006-0042-7

Zanon, M., Potrich, D., Bortot, M. & Vallortigara, G. Towards a standardization of non-symbolic numerical experiments: GeNEsIS, a flexible and user-friendly tool to generate controlled stimuli. Behav. Res. Methods, 1–12 (2021). https://doi.org/10.3758/s13428-021-01580-y

Zappia, J. V. & Rogers, L. J. Sex differences and reversal of brain asymmetry by testosterone in chickens. Behav. Brain Res. 23(3), 261–267 (1987). https://doi.org/10.1016/0166-4328(87)90026-X

